# Bacterial Outer Membrane Vesicles Induce a Transcriptional Shift in Arabidopsis Towards Immune System Activation Leading to Suppression of Pathogen Growth in Planta

**DOI:** 10.1101/2021.12.20.473005

**Authors:** Laura Chalupowicz, Gideon Mordukhovich, Nofar Assoline, Leron Katsir, Noa Sela, Ofir Bahar

## Abstract

Gram negative bacteria form spherical blebs on their cell periphery, which later dissociate and released into the surrounding environment. Previous studies have shown that these nano scale structures, derived primarily from the bacterial outer membrane and are termed outer membrane vesicles (OMVs), induce typical immune outputs in both mammals and plants. On the other hand, these same structures have been shown to promote infection and disease. To better understand the broad transcriptional change plants undergo following exposure to OMVs, we treated *Arabidopsis thaliana* (Arabidopsis) seedlings with OMVs purified from the Gram-negative plant pathogenic bacterium *Xanthomonas campestris* pv. *campestris* and performed RNA-seq analysis on OMV- and mock-treated samples at 2, 6 and 24 h post challenge. We found that the most pronounced transcriptional shift occurred in the first two time points, as was reflected by both the number of differentially expressed genes (DEGs) and the average fold change. Gene ontology enrichment analysis revealed that OMVs induce a major transcriptional shift in Arabidopsis towards immune system activation, upregulating a multitude of immune-related pathways including a variety of immune receptors and transcriptional factors. Comparing Arabidopsis response to OMVs and to single purified elicitors, revealed that while OMVs induce a similar suite of genes and pathways as single elicitors, some differential pathways activated by OMVs were detected including response to drug and apoptosis, which may indicate exposure to toxic compounds via OMV. To examine whether the observed transcriptional shift in Arabidopsis leads to an effective immune response, plants were pretreated with OMVs and then inoculated with a bacterial pathogen. OMV-mediated priming led to a significant reduction in bacterial titer in inoculated leaves two days following inoculation. Mutations in the elongation factor receptor (EFR), flagellin receptor (FLS2), or the brassinosteroid-insensitive 1–associated kinase (BAK1) receptor, did not significantly affect OMV-priming. All together these results show that OMV induce a broad transcriptional shift in Arabidopsis leading to upregulation of multiple immune pathways, and that this transcriptional change is reflected in the ability to better resist bacterial infection.

## INTRODUCTION

All plants, including agricultural crops are constantly confronted with harming microbes that thrive on plants, and on course, causing damage to the plant. An efficient defense response to such microbes depends greatly on rapid and accurate detection and identification of the invading microbe. For this purpose, plants utilize broad surveillance systems to monitor for pathogen invasion (Cook *et al*., 2015). It is speculated that the first line of the plant surveillance system, or the first cellular interface where plants and microbes interact, are the intercellular spaces, the apoplast. There, recognition of invading microbes is mediated by plasma membrane-bound, extracellularly exposed pattern recognition receptors (PRRs) (Couto & Zipfel, 2016; Boutrot & Zipfel, 2017). These membrane-bound receptors recognize microbial determinants that are widely present and conserved among many microbes and are known as, microbe- or pathogen-associated molecular patterns (MAMPs) (Ranf *et al*., 2016).

Since microbes undergo mutagenesis at a fast rate, evolutionary useful immune receptors are adapted to detect highly conserved regions of crucial microbial components that cannot be easily discarded or mutated, because of a serious fitness cost. For example, the bacterial flagellin is a crucial element in the lifestyle of many microbes including pathogens and is currently one of the best studied MAMPs (Felix *et al*., 1999; Zipfel *et al*., 2004). Perception of flagellin, or the synthetic epitope flg22 (comprised of highly conserved 22 amino acids at the N-terminus of the flagellum building block, flagellin), by the cognate plant immune receptor flagellin sensing 2 (FLS2), leads to a major transcriptional change, followed by an effective immune response that halts infection (Gómez-Gómez & Boller, 2000; Chinchilla *et al*., 2007).

Many of the known MAMPs are associated with the microbe’s cell wall. For example, fungal chitin (Fesel & Zuccaro, 2016), bacterial peptidoglycan (PG) (Gust *et al*., 2007; Erbs *et al*., 2008), bacterial lipopolysaccharides (LPS) (Dow *et al*., 2000; Silipo *et al*., 2005), flagellin (Felix *et al*., 1999; Boutrot & Zipfel, 2017), and more. Nevertheless, it is not quite clear how these cell-wall associated components are interacting with their cognate immune receptors *in planta*. Whether this occurs due to cell death and/or degradation of the cell wall, or via active release of components such as the flagellum, is a topic that has been somewhat overlooked (Bahar, 2020).

An example of active release of cell-wall fragments by Gram-negative bacteria is the release of outer membrane vesicles (OMVs), a common process in Gram-negative bacteria (Schwechheimer & Kuehn, 2015). These vesicles bleb and pinch off the outer membrane and are released to the surrounding environment of the cell. This process occurs continuously and under various environmental conditions, including during host colonization (Ionescu *et al*., 2014; Jin *et al*., 2011; Gurung *et al*., 2011). In addition to integral outer membrane molecules such as outer membrane (OM) proteins, LPS, and lipids, OMVs encapsulate periplasmic fluids, consisting of a diverse array of molecules such as proteins, cell wall degrading enzymes, polysaccharides, and nucleic acids (Kuehn & Kesty, 2005). Since OMVs are released during host colonization and since their cargo consists also of MAMPs it is tempting to speculate that they may act as carriers of immune elicitors delivering the eliciting molecules to close proximity with their cognate immune receptors (Bahar, 2020). Indeed, OMVs have been shown to induce both the mammalian and the plant immune systems when presented to their hosts (Ellis & Kuehn, 2010; Bahar *et al*., 2016; *McMillan et al*., 2020; Janda *et al*., 2021). However, while in mammalian cells the LPS component of the OMVs is a principal immune elicitor (Ellis *et al*., 2010), in plants, it is not yet known which of the OMV molecules is the prime immune elicitor.

In addition to carrying MAMPs and modulating the host immune response, OMVs were also shown to carry virulence factors, and to be involved in a multitude of processes. This includes cell-cell communication (Mashburn & Whiteley, 2005; Deatheragea & Cooksona, 2012; Raposo & Stahl, 2019), delivery of toxins to target cells (Kadurugamuwa & Beveridge, 1996; Ellis & Kuehn, 2010), biofilm formation (Schooling & Beveridge, 2006), quenching of antimicrobial compounds (Manning & Kuehn, 2011), response to stress (MacDonald & Kuehn, 2013), horizontal gene transfer (Fulsundar *et al*., 2014; Velimirov & Ranftler, 2018) and virulence (Ellis & Kuehn, 2010; Kunsmann *et al*., 2015). While most of these examples come from studies of mammalian bacterial pathogens, recent studies with plant pathogenic bacteria also support the notion that OMVs promote bacterial virulence and plant colonization. Ionescu *et al*. (Ionescu *et al*., 2014) showed that OMV production by the plant pathogen *Xylella fastidiosa* during xylem vessel colonization inhibits bacterial attachement to xylem walls, tilting the balance between the sessile and mobile forms of the pathogen towards the mobile form. This form is believed to promote cell dispersion in the xylem, leading to a faster decline of the plant (Ionescu *et al*., 2014). Two other studies have shown that virulence factors such as type II-secreted lipases/esterase and xylanase, and type III-secreted effectors, are secreted in association with OMVs (Sidhu *et al*., 2008; Chowdhury & Jagannadham, 2013; Solé *et al*., 2015) suggesting that OMV may have an important role in bacterial virulence.

The molecular complexity of OMVs, along with its dual and possibly contracting functions in the host, prompts us to study the broader transcriptional response of *Arabidopsis thaliana* (Arabidopsis) plants to OMV challenge and to test whether this transcriptional change would induce resistance, or susceptibility to subsequent bacterial infection.

## RESULTS

### RNA-seq analysis reveals a large set of Arabidopsis genes differentially expressed in response to OMV challenge

To study the transcriptional change in Arabidopsis following OMV challenged, we treated Arabidopsis seedlings with OMVs purified from the bacterial pathogen *Xanthomonas campestris* pv. *campestris* 33913 (*Xcc*) and collected plant RNA at 2, 6 and 24 h post challenge. RNA from OMV- and mock-treated samples was then sequenced and analyzed as described in Materials and Methods.

Principal correlation (Fig. 1A) and sample correlation matrix analyses (Fig. 1B) show that the biological replicates in each treatment cluster closely together, an indication of the overall transcriptional response similarity between biological replicates. The OMV treated samples at 2 and 6 h post challenge cluster together, and not with their respective mock samples, indicating that the OMV treatment had a greater effect than the time of sample collection, on the transcriptional response. The OMV treated samples at 24 h post challenge, however, cluster together with its mock treatment, suggesting that the transcriptional change 24 h post OMV challenge was less pronounced than the effect of sample collection time at this specific time point.

**Figure 1:**
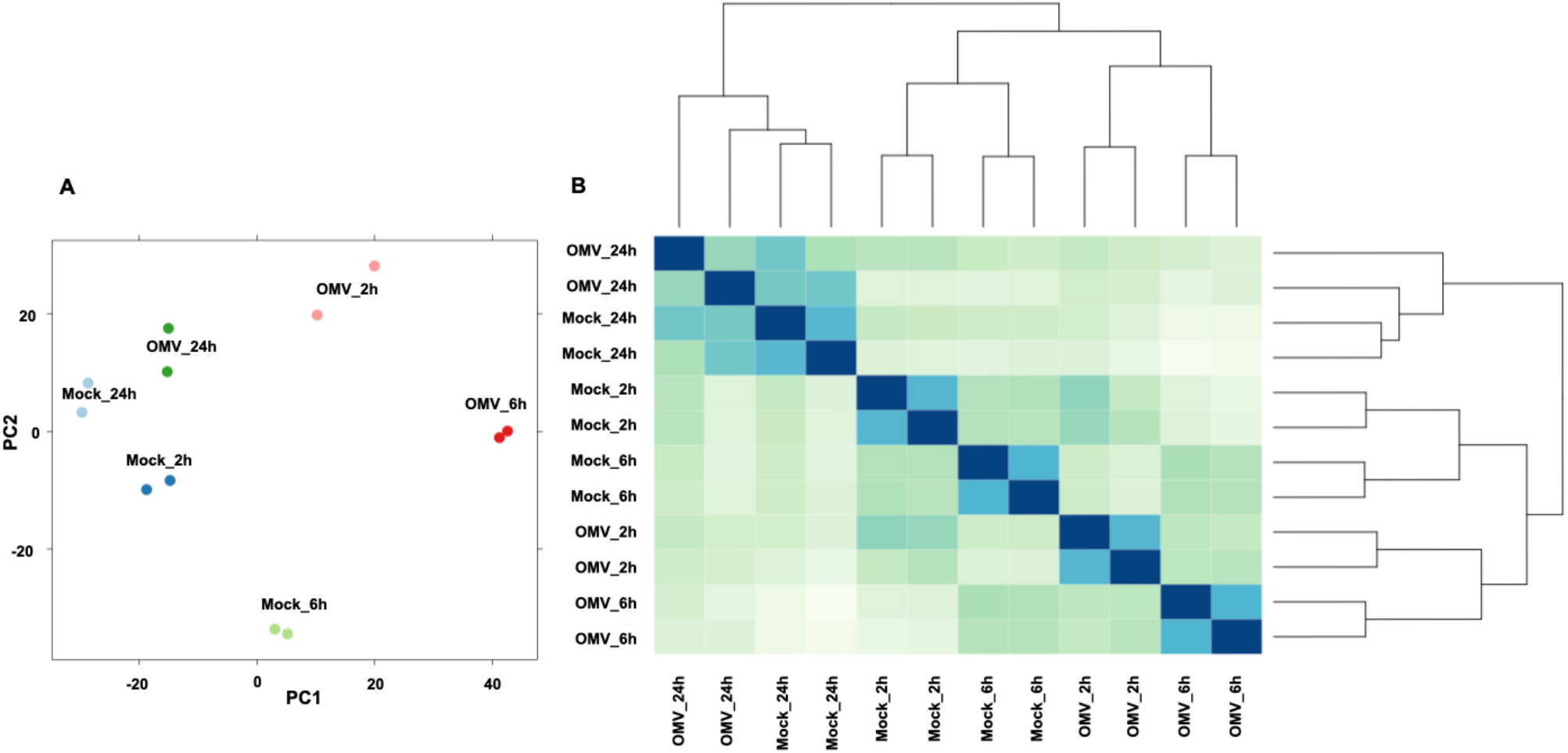
Arabidopsis transcriptional response to OMVs at 2, 6 and 24 post challenge. Principal component analysis **(A)** and sample correlation matrix **(B)** of Arabidopsis seedlings transcriptional response to OMV, or mock, at 2, 6 and 24 h post challenge.

At each of the time points tested, the transcriptome of OMV-treated Arabidopsis seedlings was compared with mock-treated seedlings and differentially expressed genes (DEGs) were extracted. In all time points combined, a total of 984 and 175 genes were found to be significantly (Log fold-change >1 or <-1, *p* value and FDR <0.05) up- or down-regulated, respectively, in response to OMV challenge (Fig. 2A; Sup. Table S1). Gene expression Log fold-change (LogFC) ranged from a maximum induction of 9.08 (AT1G26410, 6 h post challenge), which corresponds to over 500-fold change, to -5.73 (AT3G17520, 24 h post challenge). The highest number of DEGs was found at the 2 and 6 h time points, where a total of 647 and 876 DEGs, respectively, were identified (up- and down-regulated combined). At 24 h post OMV challenge, 121 DEGs were found. More than 50 % of the up-regulated genes at 2 and 6 h post OMV challenge were shared between them (Fig. 2B).

**Figure 2:**
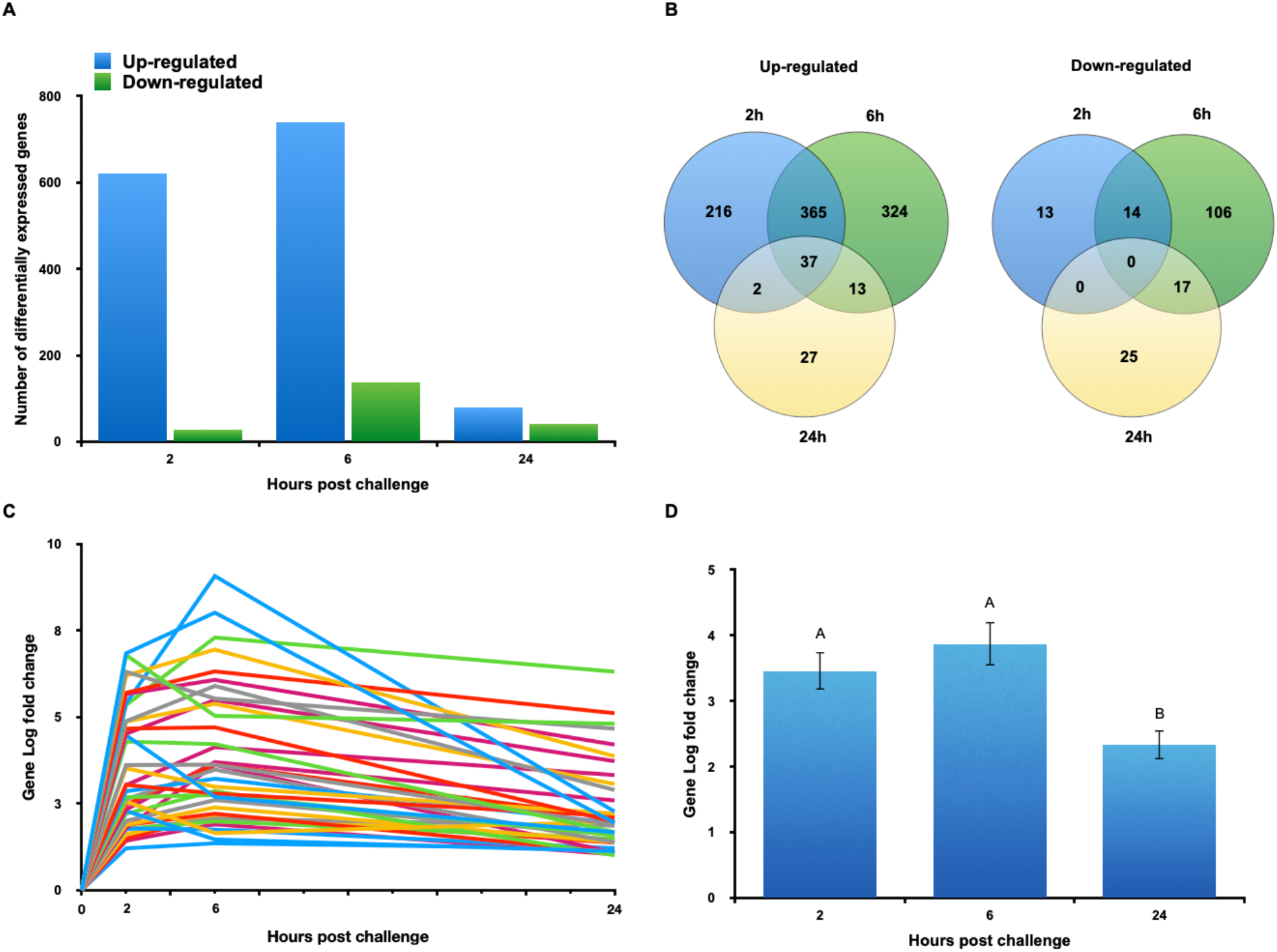
Arabidopsis differentially expressed genes in response to OMV challenge. **(A)** Total number of differentially expressed genes (DEGs) (up- or down-regulated, LogFC >1 or <-1, *p* value and FDR <0.05) at 2, 6, and 24 h post challenge. **(B)** Overlap between DEGs at different time points (left, up-regulated; right, down-regulated). **(C)** Up-regulated genes found in all three time points were plotted on a LogFC expression graph, showing gene expression over time. **(D)** LogFC average of all DEGs at different time points. Different letters indicate statistical difference at *p*<0.05 by the Tukey-Kramer HSD test.

To examine the temporal gene expression change, we extracted all the up-regulated DEGs that were found in all three time points (37 genes) and compared their fold-change throughout time (Fig. 2C). The fold-change expression of these genes was significantly different among the different time points (one-way ANOVA; *F*_2,108_ = 8.1633, *p* = 0.0005). A post hoc comparison using the Tukey Kramer HSD test indicates that gene LogFC in both the 2 h (M = 3.45, SD= 1.72) and the 6 h (M = 3.87, SD = 1.98) time points was significantly higher than the 24 h (M = 2.32, SD = 1.31) time point (*p* = 0.0145 and *p* = 0.0005, respectively) (Fig. 2D). While the mean LogFC expression at the 6 h time point was higher than that at the 2 h time point, it was not statistically different (*p* = 0.5413). Hence, when considering the number of DEGs and the overall gene LogFC at the three time points tested, we can conclude that the most significant transcriptional change occurred at 2 and 6 h post OMV challenge. To examine the validity of the RNA-seq results, the expression of 17 up-regulated genes was determined using quantitative-PCR (qPCR) with specific primers. Fourteen of the tested genes displayed the same pattern as in the RNA-seq analysis and were significantly up-regulated compared with mock. Three of the tested genes had a higher relative expression, but were not significantly different from mock by this method (Sup. Fig. S1).

### Arabidopsis responds to OMVs with a transcriptional shift towards activation of the immune system

To identify Arabidopsis pathways significantly affected by OMV challenge, we used the AgriGO web tool (Du *et al*., 2010; Tian *et al*., 2017). We identified 333 and 55 significantly (FDR<0.05) up- and down-regulated gene ontology (GO) terms, respectively, in all time points combined in response to OMV challenge. Many of the most significantly up-regulated GOs were related to plant response to biotic and abiotic factors and included the terms ‘defense response’, ‘innate immune response’ ‘response to stress’ and ‘response to biotic stimulus’ (Sup. Table S2). The AgriGO tool also identified 103 significantly up-regulated molecular functions including ‘transferase activity’, kinase activity’, ‘transcription factor activity’, ‘ion and metal ion binding’, ‘carbohydrate binding’, ‘protein binding’, ‘catalytic activity’, ‘Adenyl nucleotide binding’, ‘transmembrane receptor activity’ and more binding functions (Sup. Table S2). The cellular location of the significant terms was in different compartments of the cell including the nucleus, vacuole, and endomembrane system, but was most notably associated with the cell periphery and included ‘plasma membrane’, ‘extracellular region’, ‘cell wall’ and ‘apoplast’ ontologies.

The significantly down-regulated GOs, on the other hand, included the terms ‘toxin catabolic and metabolic process’, ‘response to water and water deprivation’, ‘lipid transport and localization’ and others (Sup. Table S2). GOs related to response to biotic stimulus were not enriched in the down-regulated genes. Significantly down-regulated molecular functions included many redox terms such as ‘oxidoreductase activity’, ‘heme binding’, ‘iron ion binding’, ‘lipid binding’, ‘oxygen binding’ and more oxygen-related functions (Sup. Table S2). Down-Regulated terms were also located to the extracellular region.

When we examined the up- and down-regulated GOs for overlap, we found that they share 32 GOs. (Sup. Table S3). Unique up-regulated GOs included, ‘defense response’, ‘response to bacterium’, ‘immune system process’, ‘innate immune system’, ‘protein phosphorylation’, ‘response to SA’, ‘response to decreased oxygen levels’, ‘cell death’, ‘camalexin biosynthesis’, and more. On the other hand, GOs related to, ‘lipid transport and binding’ and ‘iron ion binding’ were among the unique GOs in the down regulated group (Sup. Table S3).

To determine to which GO terms DEGs with the highest expression belong, we filtered the original DEGs list by selecting genes that had a LogFC higher than 4, or smaller than -4 (corresponding to 16-fold difference). We identified 117 genes that met these criteria, of which 115 were up-regulated and 2 down-regulated in all time points combined. Because of the small number of down-regulated genes with a LogFC of less than -4, no significantly repressed GOs were identified. On the other hand, 72 significantly up-regulated GOs were identified, of which the most significant ones are related to ‘response to external biotic stimulus’, ‘response to other organisms’, ‘cellular response to oxygen levels’, ‘defense response’, ‘response to stress’ and more immune related GOs (Fig. 3).

**Figure 3:**
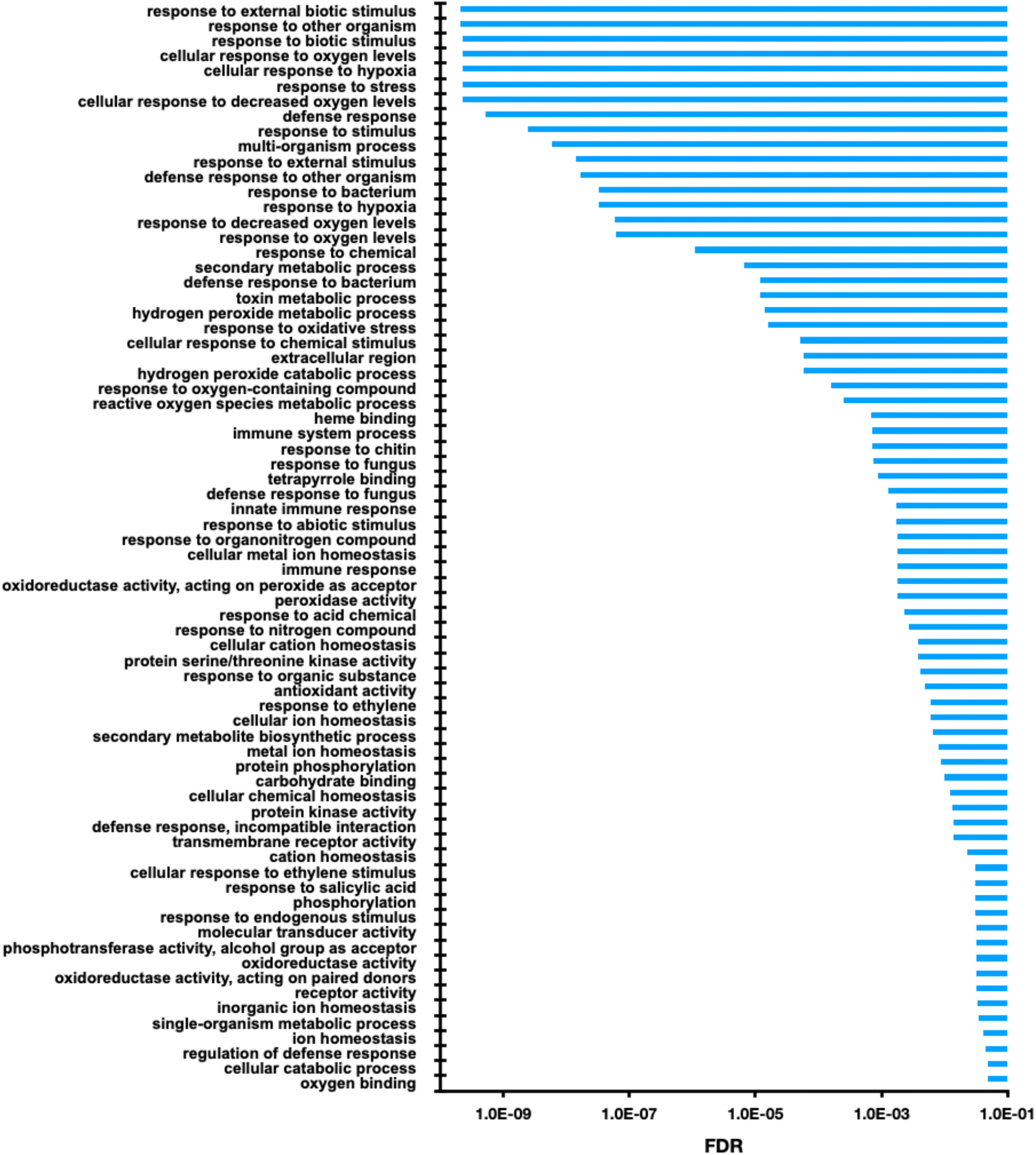
Arabidopsis gene ontology (GO) terms enriched in response to *Xcc* OMV challenge. Arabidopsis genes being highly up-regulated (LogFC of >4) in response to OMV challenge were filtered from the complete dataset of DEGs and used to identify enriched GO terms using AgriGo webtool. FDR cutoff < 0.05.

### OMV challenge led to up-regulation of immune receptors

MAMP sensing and plant response to MAMPs are largely mediated by membrane-bound PRRs that are crucial for pathogen perception and efficient mitigation of infection. PRRs are commonly classified in two groups, including receptor kinases (RKs) and receptor-like proteins (RLPs) (Boutrot & Zipfel, 2017). To identify PRRs that were differentially expressed in response to OMVs we have used previously established lists of Arabidopsis RKs (Kemmerling *et al*., 2011; Mott *et al*., 2016) and RLPs (Wang *et al*., 2008) and overlapped these lists with our DEG set. We have identified 33 and 10 up-regulated RKs and RLPs, respectively, in our OMV-challenged Arabidopsis transcriptome, in all time points combined (Table 1). Interestingly, we could not find any RKs or RLPs in our list of 175 down-regulated genes. Kemmerling *et al*. (2011) defined a list of 49 RKs, whose expression was significantly induced by MAMPs such as flg22 and NLP (necrosis and ethylene -inducing peptide 1-like protein), or pathogen treatment. We compared this list with the up-regulated RLKs from our experiment and found that 45 % of the RKs defined by Kemmerling *et al*. (2011) were also induced in response to OMVs. Among those, noteworthy are FRK1, SOBIR1, SERK4, RLK/IKU2, PSKR1, HAESA, EFR, BIR and IOS1 (Table 1). While membrane bound RKs and RLPs mediate mostly outer-cellular sensing of invading microbes, nucleotide-binding site–leucine-rich repeat (LRR) receptors (NLRs), are intracellular immune receptors. We found 7 different NLR genes up-regulated in response to OMV-challenge at 2 and 6 h post challenge, however none were found at the 24 h time point (NLR list was extracted from TAIR, 102 genes) (Table 1).

**Table 1.** OMV-induced RKs/RLPs and immune-related transcription factors in Arabidopsis seedlings.

| Gene family | Gene ID | Gene expression (LogFC) per time point (h) |  |  | Gene name |
| --- | --- | --- | --- | --- | --- |
|  |  | 2 | 6 | 24 |  |
| **RKs** |  |  |  |  |  |
|  | AT2G02220 | 1.07 |  |  | PSKR1 |
|  | AT4G28490 | 1.14 |  |  | HAESA |
|  | AT5G48380 | 1.30 |  |  | BIR |
|  | AT5G20480 | 1.12 |  |  | EFR |
|  | AT1G09970 | 1.40 | 1.03 |  | RLK7/IKU2 |
|  | AT1G51800 | 3.90 | 3.08 |  | IOS1 |
|  | AT1G51850 | 3.50 | 2.46 |  | SIF2 |
|  | AT1G74360 | 2.50 | 2.63 |  | NIRL1 |
|  | AT2G13790 | 1.60 | 1.16 |  | SERK4 |
|  | AT2G19190 | 7.11 | 5.20 |  | FRK1 |
|  | AT2G31880 | 1.62 | 1.53 |  | SOBIR1 |
|  | AT4G08850 | 1.89 | 1.63 |  | MIK2 |
|  | AT5G25930 | 2.57 | 2.06 |  | - |
|  | AT1G51790 | 3.00 | 2.02 |  | - |
|  | AT1G35710 | 2.28 | 1.46 | 1.12 | - |
|  | AT1G07560 | 1.03 |  |  | - |
|  | AT1G56140 | 1.34 |  |  | - |
|  | AT2G13800 | 1.39 |  |  | - |
|  | AT2G28960 | 1.74 | 2.06 | 1.86 | - |
|  | AT2G28970 | 1.60 | 1.23 |  | - |
|  | AT5G49780 | 1.96 | 2.31 |  | - |
|  | AT5G42440 | 1.26 |  |  | - |
|  | AT3G53590 |  | 3.16 |  | - |
|  | AT1G05700 |  | 3.97 |  | - |
|  | AT1G51820 |  | 2.33 |  | - |
|  | AT1G51860 |  | 1.53 |  | - |
|  | AT1G51890 |  | 3.52 |  | - |
|  | AT5G59680 |  | 3.53 | 1.78 | - |
|  | AT5G37450 |  | 2.24 |  | - |
|  | AT5G49770 |  | 2.41 |  | - |
|  | AT3G47090 |  | 2.05 |  | - |
|  | AT5G44700 |  | 1.58 |  | GSO2 |
|  | AT3G09010 |  | 1.94 |  | - |
| **RLPs** |  |  |  |  |  |
|  | AT2G32680 | 3.68 |  | 2.79 | RLP23 |
|  | AT2G25470 | 3.44 | 2.41 |  | - |
|  | AT3G23120 | 2.19 |  |  | RLP38 |
|  | AT1G71400 | 1.68 | 1.48 |  | RLP12 |
|  | AT3G05660 | 1.15 |  |  | RLP33 |
|  | AT1G47890 |  | 3.92 |  | RLP7 |
|  | AT3G28890 |  | 2.50 |  | RLP43 |
|  | AT5G25910 |  | 2.04 |  | RLP52 |
|  | AT3G11080 |  | 1.96 |  | RLP35 |
|  | AT3G05360 |  | 1.63 |  | RLP30 |
| **NLRs** |  |  |  |  |  |
|  | AT4G14370 | 2.49 | 2.45 |  | - |
|  | AT5G45000 | 2.00 | 1.99 |  | - |
|  | AT5G45240 | 1.74 | 1.60 |  | - |
|  | AT5G45220 | 1.03 |  |  | - |
|  | AT5G41750 |  | 2.60 |  | - |
|  | AT2G17050 |  | 1.73 |  | - |
|  | AT5G41740 |  | 1.33 |  | - |
| **WRKY** |  |  |  |  |  |
|  | AT5G24110 | 4.65 | 6.24 |  | WRKY30 |
|  | AT1G66600 | 3.92 | 6.11 |  | WRKY63 |
|  | AT4G23810 | 3.10 | 3.60 |  | WRKY53 |
|  | AT2G38470 | 2.90 | 2.16 |  | WRKY33 |
|  | AT1G80840 | 2.55 | 1.64 |  | WRKY40 |
|  | AT4G22070 | 2.18 | 2.13 |  | - |
|  | AT5G15130 | 1.85 | 1.68 |  | - |
|  | AT2G23320 | 1.46 |  |  | WRKY15 |
|  | AT4G18170 | 1.36 |  |  | WRKY28 |
|  | AT4G31550 | 1.06 |  |  | - |
|  | AT4G01250 | 1.02 |  |  | WRKY22 |
|  | AT2G30250 | 1.02 |  |  | WRKY25 |
|  | AT1G62300 | 1.44 | 1.31 |  | WRKY6 |
|  | AT2G40740 |  | 4.37 |  | - |
|  | AT5G22570 |  | 4.29 |  | WRKY38 |
|  | AT5G01900 |  | 3.80 |  | WRKY62 |
|  | AT1G29860 |  | 2.99 |  | WRKY71 |
|  | AT2G40750 |  | 2.42 |  | WRKY54 |
|  | AT5G13080 |  | 2.05 |  | WRKY75 |
|  | AT5G49520 |  | 1.24 |  | WRKY48 |
| **MYB** |  |  |  |  |  |
|  | AT3G23250 | 4.18 | 5.11 |  | MYB15 |
|  | AT4G37780 | 3.56 |  |  | MYB87 |
|  | AT1G18570 | 3.53 | 2.28 |  | MYB51 |
|  | AT1G74080 |  | 2.90 |  | MYB122 |
|  | AT4G12350 |  | 2.36 |  | MYB42 |
|  | AT5G07690 | -1.12 | -2.05 |  | MYB29 |
|  | AT4G05100 |  | -1.58 |  | MYB74 |
|  | AT3G24310 |  | -1.85 |  | MYB305 |
|  | AT3G30210 |  |  | -3.87 | MYB121 |
| **AP2/ERF** |  |  |  |  |  |
|  | AT1G71520 | 4.11 |  |  | - |
|  | AT3G23240 | 4.05 |  |  | - |
|  | AT5G64750 | 3.26 |  |  | ABR1 |
|  | AT5G61890 | 3.17 | 4.04 |  | ERF114 |
|  | AT5G47230 | 2.69 | 1.97 |  | ERF102 |
|  | AT1G28370 | 2.54 |  |  | - |
|  | AT4G17490 | 2.47 | 1.45 |  | ERF103 |
|  | AT5G51190 | 2.20 | 1.26 |  | ERF105 |
|  | AT5G47220 | 2.06 |  |  | ERF2 |
|  | AT2G33710 | 1.74 |  |  | - |
|  | AT4G17500 | 1.57 | 1.05 |  | ERF100 |
|  | AT3G50260 | 1.51 | 1.00 |  | - |
|  | AT1G72360 | 1.48 |  |  | ERF73 |
|  | AT3G25730 | 1.28 |  |  | - |
|  | AT5G61600 | 1.14 |  |  | ERF104 |
|  | AT5G13330 | 1.07 | 1.07 |  | - |
| **bHLH** |  |  |  |  |  |
|  | AT5G56960 | 4.82 |  |  | - |
|  | AT2G43140 | 1.36 | 1.78 |  | BHLH129 |
|  | AT3G56980 |  |  | 2.57 | BHLH39 |
|  | AT4G28790 |  | -1.35 |  | - |
|  | AT1G51140 |  | -1.44 |  | BHLH3 |
|  | AT1G71200 |  | -1.53 | -2.55 | BHLH160 |
|  | AT4G29930 |  | -2.19 | -1.38 | - |
| **C2H2** | AT2G37430 |  | 3.9 |  | ZAT11 |
|  | AT3G49930 |  | -1.40 |  | - |
| **bZIP** | AT1G42990 | 1.04 |  |  | BZIP60 |

### OMVs induce the expression of multiple WRKY transcription factors

WRKY transcription factors (TFs) are central to the plant immune system, participating in both MAMP-triggered immunity (MTI) and effector-triggered immunity (ETI) responses (Rushton *et al*., 2010; Birkenbihl *et al*., 2018). OMV challenge led to up-regulation of 20 different WRKY TFs (list extracted from TAIR, 70 genes) at 2 and 6 h post challenge, however none were found at the 24 h time point (Table 1). Interestingly, we could not detect any WRKY TFs in our down-regulated gene set in any of the time-points tested. Another family of TFs affected by OMV challenge are MYB domain-containing proteins (Tsuda & Somssich, 2015), known to be involved in multiple processes including biotic and abiotic stresses (Ambawat *et al*., 2013). Overall, 9 different MYB TFs (list extracted from TAIR, 211 genes) were differentially-expressed in response to OMV challenge, 5 up-regulated and 4 down-regulated (Table 1). Other major transcriptional factors involved in plant immunity were also differentially-expressed in response to OMV challenge. Sixteen members of AP2/ERF family were found to be up-regulated at the 2 and 6 h time points combined, and seven differentially-expressed bHLH TFs were also identified. Additional types of differentially-expressed TFs are listed in Table 1.

### Comparing Arabidopsis transcriptional response to OMV with response to single MAMPs

OMVs are molecularly much more complex than single, synthetic MAMPs/DAMPs such as flg22, elf18/26, chitin, oligogalacturonides (OGs), or others. To learn about the differences in Arabidopsis response when challenged with a single, purified elicitor versus a natural and molecularly complex eliciting structure – OMVs, we compared our RNA-seq data with existing transcriptomic data of Arabidopsis response to known MAMPs including flg22, (Denoux *et al*., 2008) elf26 (Zipfel *et al*., 2006), PGN (Willmann *et al*., 2011), OGs (Davidsson *et al*., 2017) and LPS (Livaja *et al*., 2008). Enriched GOs were extracted from the above-mentioned datasets as describe above (Sup. Table S4), and they were compared with GOs enriched following OMV challenge. Generally, Arabidopsis GO terms induced by OMVs were similar to those induced by single, proteinaceous and non-proteinaceous MAMPs, sharing 56, 51 and 47 % of OMV-induced GOs with GOs induced by flg22, elf26 and PGN, respectively. On the other hand, a lower overlap in induced GOs was seen with LPS and OGs, sharing 24 and 33 %, respectively, with OMV-induced GOs (Fig. 4A). Notably, the pathogenesis-related 1 (PR1) gene (At2g14610), a hallmark of LPS-induced immune responses (Silipo *et al*., 2005, 2008), was absent from the OMV-induced gene list in all the time points tested. Comparing OMV induced GOs with GOs induced by other MAMPs tested here, yielded a set of 41 GOs found only in the OMV treatment (Fig. 4B). This list included GOs related to ‘apoptosis’, ‘response to drug’, ‘drug transport and ‘multi-drug transport’, ‘lipase activity’, and more (Sup. Table S4, denoted by asterisks and bold font).

**Figure 4:**
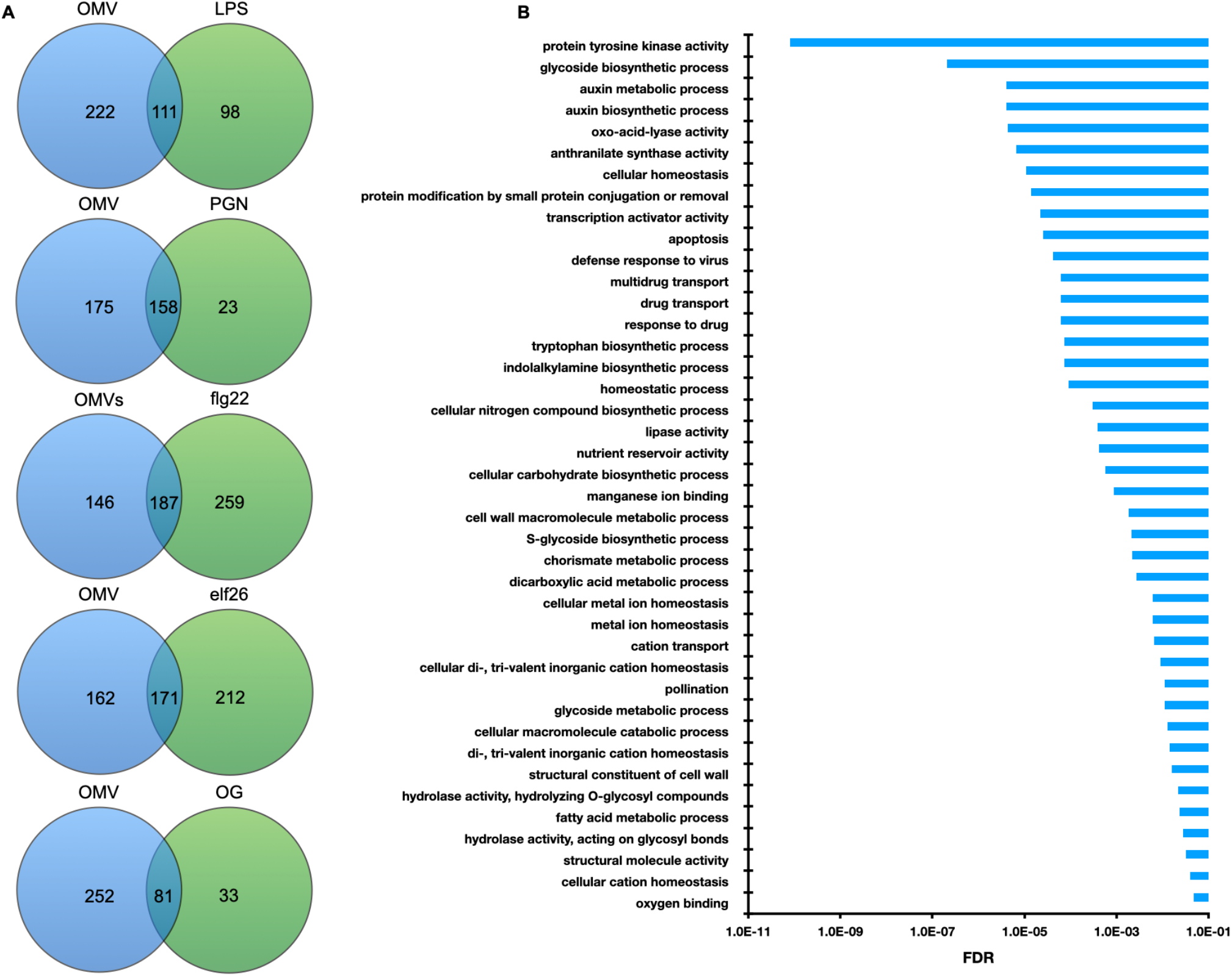
Comparison of enriched Gene ontology (GO) terms in response to OMV and to different single purified MAMPs. Arabidopsis expression datasets in response to MAMP challenge (elf26, flg22, OGs, PGN and LPS; see Materials and Methods section for references) were used to extract enriched GOs using the AgriGo web-tool. Enriched GO sets of each MAMP were compared with the enriched GO list in response to OMVs using Venny (**A**, and Sup. Table S4). GO terms enriched only in the OMV datasets are shown in (**B**) sorted by their FDR value.

### OMVs induce Arabidopsis resistance to bacterial infection

Here and previously (Bahar *et al*., 2016), we have provided evidence demonstrating that the Arabidopsis immune system is induced by OMV challenge. To examine whether this OMV-mediated immune induction is translated into an effective immune response, we used an *in planta* bacterial growth test (Zipfel *et al*., 2004) in which Arabidopsis plants are pretreated with OMVs followed by bacterial inoculation. A significant decrease of more than 10-fold in *Pseudomonas syringae* pv. *tomato* DC3000 (*Pst*) CFU/g leaf was observed in both OMV- and flg22-pretreated plants compared with mock pretreatment, two days after inoculation (Fig. 5A). *In vitro* growth of *Pst* was not negatively affected by addition of OMVs to the medium, suggesting that the reduced *Pst* growth *in planta* is related to the priming effect OMV had on the plant, and is not a direct effect of OMVs on the bacteria (Sup. Fig. S2). To test whether the OMV-induced resistance to *Pst* is mediated by FLS2, EFR or BAK1 we repeated this experiment with Col-0, *bak1-5* mutant and the double mutant line *fls2 efr1*. With both mutant lines, OMV pretreatment resulted in a significant reduction in *Pst* CFU/g leaf compared with mock treated plants (Fig. 5B-C). As expected, *fls2 efr1* and *bak1-5* mutant lines treated with flg22 had similar *Pst* titers as the untreated plants as they are known to be irresponsive to flg22. To compare the relative reduction in pathogen titer in Col-0 versus the immune receptor mutant lines *fls2 efr1* and *bak1-5* in primed plants, we calculated the difference in *Pst* titer in OMV- and mock-treated plants in three independent experiments (Sup. Fig. S3). The average reduction in *Pst* titer in OMV-pretreated *bak1-5* plants was smaller than that observed in Col-0 plants (0.89 Vs.1.14 Log CFU/gr leaf reduction for *bak1-5* and Col-0, respectively, one-way ANOVA; *F*_2,4_ = 4.3781, *p* = 0.0523). We did not see a similar reduction with the *fls2 efr1* mutant line (1.26 Vs. 1.32 Log CFU reduction for *fls2 efr1* and Col-0, respectively, one-way ANOVA: *F*_1,4_ = 0.0352, *p* = 0.5698) (Fig. 5D-E).

**Figure 5:**
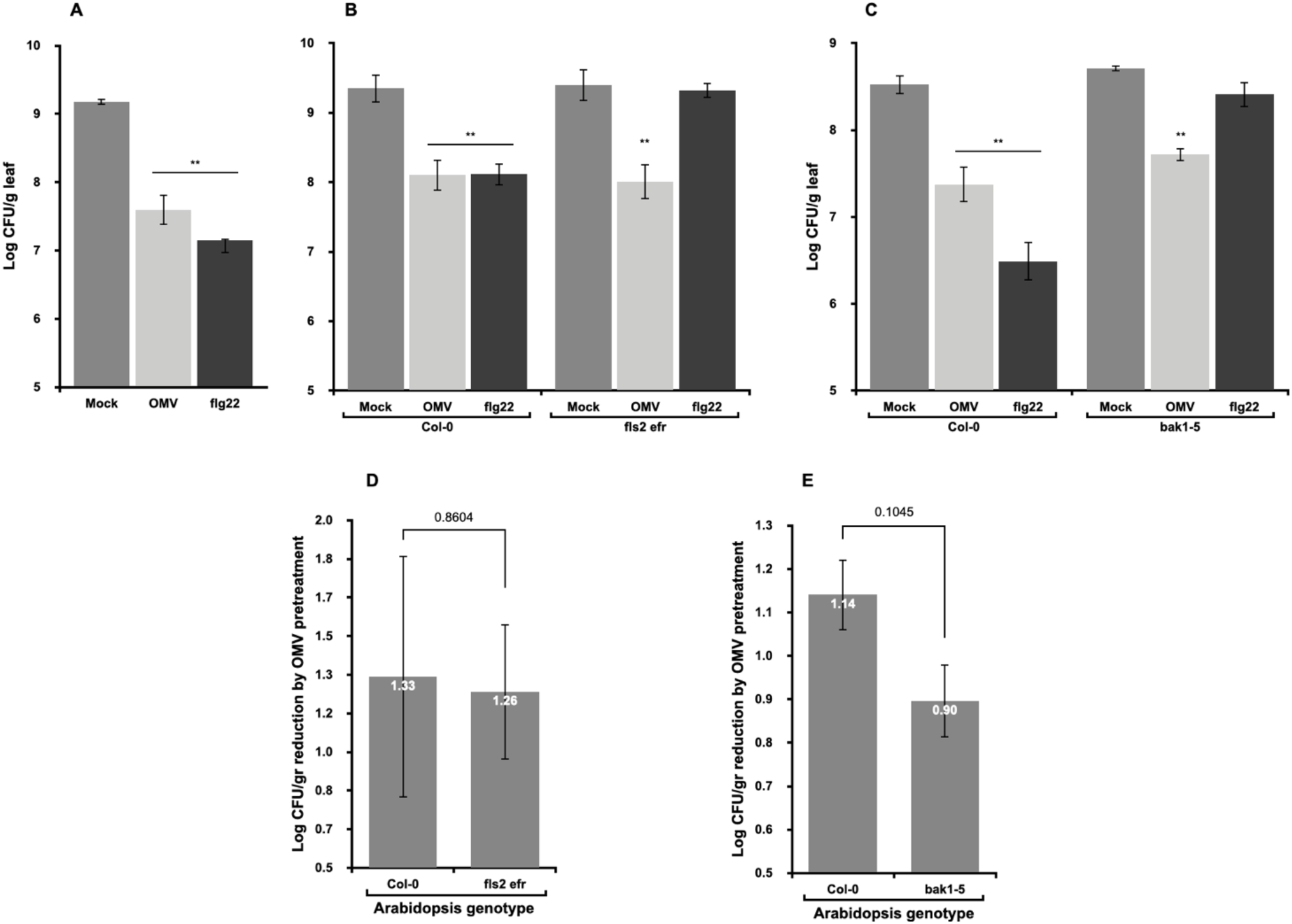
Pretreating Arabidopsis leaves with OMVs induce resistance to subsequent bacterial infection. Col-0 plants **(A)** were pretreated with OMVs, water (mock) or flg22 as controls, and 24 h later inoculated with a 10^5^ CFU/mL suspension of *Pst* DC3000 using needleless syringe infiltration. *Pst* DC3000 cell titer in the inoculated leaves was determined 48 h after inoculation by serial dilution platings. Arabidopsis Col-0 and *fls2 efr* **(B)**, or *bak1-5* **(C)** plants were tested in a similar experiment as described in (A). The mean Log *Pst* DC3000 CFU/gr reduction following OMV pretreatment (compared with untreated plants) in Col-0 and *fls2 efr* **(D)**, and Col-0 and *bak1-5* **(E)**, was compared. Each bar represents the mean Log *Pst* DC3000 CFU/gr reduction from three independent experiments (data of the independent experiments is presented in Sup. Fig. S3). Differences were not statistically significant (Two-tail student t-test. *p* values are indicated above the graph bars). Experiment A, B, and C were conducted at least three times with similar results (3 plants/replicates per treatment in each experiment). Asterisks (**) indicates significant difference compared with mock (Dunnet’s test *p*<0.001).

## DISCUSSION

Bacterial outer membrane vesicles (OMVs) are complex nanostructures originating from the bacterial outer membrane and are composed of hundreds of proteins, as well as other cell wall components. It was previously shown that Arabidopsis plants respond to OMV challenge by activating typical immune responses such as ROS burst, immune marker gene expression and medium alkalization (Bahar *et al*., 2016). In this study we examined the broader transcriptional response of Arabidopsis to bacterial OMVs, and its effect on subsequent infection.

The overarching conclusion coming from the RNA-seq data analyses performed in this study, is that the Arabidopsis immune system is primed following the exposure to *Xanthomonas campestris* pv. *campestris* (*Xcc*) OMVs. This conclusion is supported by different and complementing analyses. **First**, gene ontology (GO) enrichment in plants exposed to *Xcc* OMVs clearly show that the OMVs are perceived by Arabidopsis as stressors. The cellular location of the plant response was associated primarily with the cell periphery suggesting outer cellular perception of the challenging material, OMVs. This provides further support to the notion that OMVs, and their constituents are sensed by extracellular receptors, similarly to many known MAMPs. **Secondly**, we noticed a large suite of RKs and RLPs upregulated in response to OMV challenge. Many of these receptors are known to mediate pathogen perception or were previously shown to associate with plant immune response. Interestingly, FLG22-induced receptor-like kinase 1 (FRK1) was the most highly induced receptor in both 2 and 6 h post challenge with a LogFC of 7.11 and 5.2, respectively, while the average LogFC of all RKs was 2.11 and 2.13 at 2 and 6 h, respectively. This is interesting since we could not detect flagellin in our *Xcc* OMV proteomics analysis (data not shown). It is known that FRK1 is also induced by other immune elicitors, but it is intriguing why its expression is so much higher than the rest of the RKs up-regulated here. Elongation factor receptor (EFR) expression on the other hand, was only significantly up-regulated at the 2 h time point, and had a LogFC of 1.12, even though EF-Tu is found in *Xcc* OMVs (Bahar *et al*. 2016). We also found a few NLR genes up-regulated in response to OMV challenge, half of which are annotated as disease resistance proteins, yet their function in plant immunity has not been described. While we do not hypothesize NLRs are directly involved in OMV perception, they could be induced downstream to RK/RLPs sensing of OMV molecules as was also observed in response to purified MAMPs such as flg22, elf18 and LPS (Livaja *et al*., 2008; Denoux *et al*., 2008; Zipfel *et al*., 2006). **Thirdly**, many immune-related transcription factors members of the WRKY, MYB and other families were significantly upregulated by OMVs (Bjornson *et al*., 2021).

In our study, the main transcriptional change in response to OMV occurred at the first two time points (2 and 6 h post challenge). This was illustrated by both a significantly larger number of differentially expressed genes (DEGs) and a significantly higher Log fold-change (LogFC) in the expression level of genes in the 2 and 6 h time points versus the 24 h post challenge time point. Nevertheless, out of a total of 121 DEGs found in the 24 h time point, almost half (52) were not found in the 2 or 6 h time points. This suggests that half of the DEGs at the 24 h time point are late-regulated genes, whose expression was up- or down-regulated later than 6 h post challenge. Indeed, Arabidopsis genes with different expression dynamics following elicitor challenged were previously identified (Bjornson *et al*., 2021).

The rapid and mostly transient gene expression pattern we have seen here is in accordance with other studies that have tested the temporal response of Arabidopsis to MAMPs. For example, Denoux *et al*., (2008) and Bjornson *et al*., (2021) have shown that the transcriptional change in Arabidopsis in response to various MAMPs occurs within minutes to hours, and that this change is transient, and in most cases, DEGs are back to base levels ∼24 h following plant challenge. Unlike bilateral interactions between a plant and a pathogen, where each of the partners responds to the changing environment and the interaction is dynamic and ongoing, when challenging with a non-living sample, such as synthetic or purified MAMPs as well as with OMVs, the interaction is unilateral in principle. Hence, it can be expected that the response to such non-living elicitors, at least on a transcriptional level, would be transient and not sustained over days.

Intensive research in the past three decades have revealed multiple plant immune receptors responsible for microbe recognition. Many of these receptors have the capacity to detect single microbial features and are being studied in detail to better understand pathogen perception, immune system signaling and response of model and crop plants. In real-life, plants are simultaneously exposed to multiple microbial features from different sources, which interact with their immune system. Therefore, we were interested to examine the differences in the transcriptional response of Arabidopsis to a singular purified MAMPs versus OMVs, which represent a more natural and complex microbial structure. Additionally, while synthetic MAMPs are not hypothesized to have an activity *in planta*, other than being detected by the immune system, OMVs carry virulence factors, degradative enzymes, toxins and other biomolecules that could have a functional role while in the plant, and therefore it was interesting to test if OMV challenge induces unique GOs that are not induced by synthetic MAMPs.

For our transcriptomic comparisons, we collected data from studies with similar experimental settings as possible to ours, i.e., similar time points, challenging plants and not cell cultures or protoplasts, and so on. A significant overlap in GO enrichment was seen in Arabidopsis response to OMV and Arabidopsis response to the MAMPs flg22, elf26 and PGN. This is not unexpected as it is known that many of the defense pathways activated upon pathogen sensing are similar, regardless of the specific elicitor and its source (Zipfel *et al*., 2006; Bjornson *et al*., 2021). Nevertheless, some unique GOs were found to be upregulated by OMVs and not by the other MAMPs we have surveyed. Among these are GOs related to cells wall degradation such as ‘lipase activity’ and ‘hydrolase activity acting on glycosyl bonds’, which may indicate that the plant defense system is targeting OMV degradation. Interestingly, three GOs related to drug transport were also found to be uniquely upregulated by OMV challenge. This may indicate that during OMV challenge, plants are faced with toxic compounds being delivered into their cells, perhaps by OMV-mediated delivery.

Unlike Arabidopsis response to immune eliciting peptides and PGN, we observed relatively little overlap between Arabidopsis response to OMVs and to OGs and LPS. This little overlap, especially with LPS, is somewhat surprising, considering that in mammalian cells, LPS are well acknowledged as potent inducers of host immune response to OMVs (Ellis *et al*., 2010). Furthermore, the fact we could not find upregulation of the LPS immune hallmark PR1 (Silipo *et al*., 2005), could suggests that LPS is not a major elicitor in bacterial OMVs. This, however, remains to be more thoroughly examined.

OMVs were shown to contribute to bacterial colonization of both mammalian and plant hosts and in some instances to bacterial virulence. On the other hand, OMVs activates the host immune system hence, acting as a double-edged sword, promoting bacterial survival and virulence on the one hand, and feeding the host surveilling system and activating host immunity on the other (McMillan & Kuehn, 2021). It was therefore intriguing to test the consequence of OMV pretreatment on subsequent bacterial infection. Our priming assays showed that OMV challenge led to a significant inhibition of *Pseudomonas syringae* pv. *tomato* DC3000 (*Pst*) growth *in planta*, similarly to the priming effect seen with synthetic MAMPs (Jung *et al*., 2009). Hence, in this instance, the pre-administration of OMVs to the inoculated tissue did not promote bacterial colonization, but rather primed the plants to induce an effective immune response that suppressed pathogen growth. This result is in line with our transcriptional data and with our previous study, supporting the notion that OMVs induce a robust and effective immune response in Arabidopsis. Two recent studies have also shown that Arabidopsis pretreatment with OMVs from a pathogenic or a commensal *Pseudomonas* species, suppressed subsequent *Pst* infection (McMillan *et al*., 2021; Janda *et al*., 2021). These results cumulatively indicate that under the conditions tested, OMV infiltration do not facilitate pathogen infection. Differently from the results mentioned above, OMV release in grapevines by the xylem-limited pathogen *Xylella fastidiosa* was shown to promote bacterial dissemination and virulence in grapevines (Ionescu *et al*., 2014). Several critical differences however can be noted among these patho-systems: first, it is not at all clear whether OMV released in grapevine xylem induces immunity or not, and secondly, *X. fastidiosa* is a xylem-limited pathogen and therefore the contribution of OMV to its virulence may involve xylem-specific mechanisms, such as affecting bacterial attachment, as was suggested by the authors. It is therefore possible that OMVs play different roles in different patho-system, depending on the lifestyle of the pathogen, the tissues it infects and the ability of the host plant to sense and respond to OMVs.

In a previous study we have shown that multiple immune receptor mutants maintain WT responsiveness to *Xcc* OMVs. These mutants included known PRRs recognizing either proteinaceous (FLS2, EFR, RLPReMAX) or non-proteinaceous MAMPs (LYM1/LYM3) (Bahar *et al*., 2016). Interestingly, Janda *et al*. (2021) reported that when the FLS2 receptor mutant line of Arabidopsis (*fls2*) was challenged with OMVs from *Pst*, FRK1 expression was unchanged and was similar to mock-treated plants, suggesting that FLS2 mediates the response to *Pst* OMVs. It is possible that flagellin is more abundant in *Pst* OMV preparations than it is in *Xcc* 33913 OMVs, and therefore the removal of the flagellin receptor had a more pronounced effect on plant response to *Pst* than to *Xcc* OMVs. Additionally, it is important to consider that the assay used by Janda *et al*., (2021) was measuring FRK1 expression only, and therefore may not provide a complete view of the *fls2* KO response to OMVs. It has been shown that different assays to test plant immune induction may yield different results, leading to allegedly contradicting conclusions. For example, we have shown that in a leaf-disc ROS burst assay, the response of Arabidopsis to OMVs was dependent on the EFR receptor, however, in immune marker gene expression assay with Arabidopsis seedlings, the *efr1* mutant line was as responsive to OMVs as the WT. More, McMillan *et al*. (2021) showed that different physical treatments applied to OMVs abolished certain activities such as seedling growth inhibition. However, it did not alter others immune outputs such as plant priming. Hence, it is important to combine a variety of assays to test different immune outputs to obtain as broad as possible view on the plant immune response to a given elicitor.

To further examine the involvement of some of the known PRRs and co-receptors, we have tested the *fls2 efr* and the *bak1-5* mutant lines using the plant priming assay. Our results show that the Arabidopsis double mutant line *fls2 efr* was similarly primed by *Xcc* OMV as were WT plants, supporting the notion that MAMPs other than flagellin and EF-Tu are also present in *Xcc* 33913 OMVs. Based on immune marker gene expression assays, we previously suggested that the BAK1 co-receptor is involved in OMV perception and/or response (Bahar *et al*., 2016). In this study, we revisited this suggestion using the priming assay. Here, the *bak1-5* mutant line was primed by OMV pretreatment, but to a slightly lesser extent than WT Col-0 plants. While this margin was not statistically significant it was greater than that seen with the double *fls2 efr* mutant line. This result is also in line with a recent study that showed that the *bak1-4* mutant line was responsive to OMV as WT plants in immune priming experiments (Tran *et al*., 2021). Overall, this may suggest that while BAK1 is involved in OMV perception, other immune perception and signaling pathways are primed by OMV, leading to an effective immune response and suppression of pathogen growth.

The involvement of the co-receptors BAK1 and SOBIR1 in response to OMVs (Bahar *et al*., 2016) led us to assume that multiple immune receptors, likely PRRs, are involved in OMV perception. However, a recent study suggested that plant immune activation by OMVs may be MAMP-independent and results from physico-chemical changes in the plant plasma membrane induced by OMV integration (Tran *et al*., 2021). This is an intriguing hypothesis that remains to be further addressed. Interestingly, McMillan *et al*., (2021) reported that OMVs treated with proteinase K retained their immune priming capacity, indicating that this activity may be independent of the OMV proteinaceous cargo. While this result may support the MAMP-independent immune activation hypothesis of Tran *et al*., (2021), other, non-proteinaceous MAMPs present in OMVs such as LPS and PGN may activate MTI (Bahar *et al*., 2016; McMillan *et al*., 2021). Additionally, proteinase K-treated OMV retained their ability to induce seedling growth inhibition, indicating that growth inhibition is dependent on the proteinaceous cargo of OMVs (McMillan *et al*., 2021). All together, these results further emphasize the complexity of plant response to OMVs and the importance of using a variety of outputs to test the involvement of a particular elicitor in specific pathways.

In summary, in 2021 alone, four independent studies including this one (that have likely taken place simultaneously), reported that bacterial OMVs modulate the plant immune system and induce an effective response against pathogen infection (Janda *et al*., 2021; McMillan *et al*,. 2021; Tran *et al*., 2021). These exciting results position OMVs as a new and important player in plant-microbe interactions, where there is still much to be learned. In this study, we provide a broader view on the transcriptional response of Arabidopsis to *Xcc* OMV. Complemental research approaches would be needed to further dissect how plants perceive OMVs and what are the components and mechanisms involved in this process.

## MATERIALS AND METHODS

### Plant material and growth conditions

*Arabidopsis thaliana* (Arabidopsis) wild type Col-0 line as well as the following mutant lines: *bak1-5* (Schwessinger *et al*., 2011) and *fls2 efr1* (Nekrasov *et al*., 2009) were used in this study. Arabidopsis seeds were surface sterilized and sown on Murashige and Skoog (MS) agar plates as described (Bahar *et al*., 2016). Plates were kept in the dark at 4°C for 2-4 days and then moved to 22°C with 16 h photoperiod for 5-8 days. Germinated seedlings of similar size were transferred into 24-well plates (two seedlings per well) containing 1 mL of MS medium with 1% (w:v) of sucrose (Duchefa Biochemie) and grown for another 8-10 days at the same conditions before challenged with elicitor as described below.

For priming assays, seeds of Arabidopsis wild type Col-0 line and mutant lines (*fls2 efr1* and *bak1-5*) were germinated as described above and then transplanted into 7×7×6 cm pots (1 seedling /pot) containing mix soil Green #7611 (Evenari, Ashdod, Israel) and grown at a 9 h photoperiod at 22°C. Plants were irrigated twice a week and fertilized using an NPK mix (6:2:4, Deshen Gat, Israel) once a week.

### Bacterial outer membrane vesicles purification

Glycerol stocks of *Xanthomonas campestris* pv. *campestris* (*Xcc*) 33913 were streaked on Nutrient Agar (Difco, NA, Becton, Dickinson and Company) plates and grown for 2-5 days at 28°C. Single colonies were collected and used to inoculate a 3-mL YEB (yeast extract broth) starter containing 10 µg/mL cephalexin hydrate (Cp, Sigma-Aldrich). Starters were grown overnight at 28°C with 185-200 rpm shaking and then used to inoculate 500 mL of PSB (peptone sucrose broth) medium with antibiotics (as described above) in 2-L Flasks at a ratio of ∼1:1000 (v:v). Cultures were grown as describe above to an OD_600_ of 0.6-0.8 and then bacterial cells were spun down and OMVs were extracted from the supernatant as described in (Mordukhovich & Bahar, 2017). The crude OMV preparation was then subjected to Optiprep gradient centrifugation to obtain purified OMVs, as described (Bahar *et al*., 2016; Mordukhovich & Bahar, 2017). Purified OMVs were kept at 4°C for up to 7 days until further use.

### Arabidopsis seedling challenge with OMVs

To examine the transcriptional response of Arabidopsis to OMV challenge, Col-0 seedlings grown in 24-well plates as described above were used. The day before OMV challenge, MS medium was withdrawn from plates and replaced with 250 µL of sterile dH_2_O and plates were left on the bench for overnight. The morning after, 20 µL of purified OMVs (30 µg per mL), or sterile dH_2_O as mock, were added to each well. Seedlings were collected 2, 6, and 24 h after challenge, blotted dry on paper and snap frozen with liquid nitrogen in 2-mL Eppendorf Safe-Lock tubes (Hamburg, Germany). At each time point, four OMV-treated and four mock-treated wells were collected, representing four biological replicates for each treatment at each time point.

### RNA purification

RNA was extracted from Arabidopsis seedlings using the TRIzol reagent (Invitrogen) according to the manufacturer instructions. RNA was further purified by using the Turbo DNA-free Kit (Ambion, Thermo Fisher Scientific), and the RNA Clean-Up and Concentration Kit (Norgen Biotek) according to the manufacturer’s instructions. Purified RNA samples were subjected to concentration and quality analyses using a TapeStation 2200 machine (Agilent Technologies), RNA Screen Tape and RNA Screen Tape Sample Buffer (Agilent Technologies), according to the manufacturer instructions and then kept at –80°C until used.

### RNA library construction and sequencing

For each treatment and time point, two samples showing the highest purity were selected for analyses. TruSeq mRNA libraries (Illumina) with PolyA capture were prepared from the selected RNA samples at the Crown Institute of Genomics at the Weizmann Institute of Science (Rehovot, Israel). Each sample was tagged, and a pool was prepared from the samples. This pool was then loaded inside two NGS lanes, and run in Illumina HiSeq sequencing machine, at high output run mode, single read (SR) 60 (v4).

### Sequencing reads initial processing

The raw sequence reads were cleaned with Trimmomatic software v 0.36 (Bolger *et al*., 2014), removing low quality reads and remaining adapter sequences. The clean reads were mapped to the reference Arabidopsis TAIR10 reference genome (Lamesch *et al*., 2012) using bowtie2 (Langmead *et al*., 2012) and quantification of genes expression was done using RSEM (Li & Dewey 2011).

Principal component analysis and sample correlation matrix were calculated with the function cor() and precomp(), respectively, of the R base package version 3.6.1. DEGs were determined using the DESeq2 tool (Love *et al*., 2014). The FDR (false discovery rate) cutoff chosen was FDR < 0.05. The LogFC (Log of the fold change) cutoff for the up-regulated and the down-regulated genes, was > 1 and < -1, respectively. Venn diagrams were built with the use of Venny 2.1 (http://bioinfogp.cnb.csic.es/tools/venny/).

### Gene ontologies and gene descriptions

Gene Ontologies (GO) were retrieved by using TAIR’s GO Annotations (http://www.arabidopsis.org/tools/bulk/go/index.jsp). Gene descriptions, according to the gene models, were retrieved by using TAIR’s gene description search (http://www.arabidopsis.org/tools/bulk/genes/). GO enrichment was calculated using the AgriGO web tool (http://bioinfo.cau.edu.cn/agriGO/index.php) using Arabidopsis genome locus (TAIR10) as reference.

### Quantitative-PCR and RNA-seq results validation

To validate RNA-seq data, RNA samples of mock- and OMV-challenged seedlings were used for cDNA synthesis, followed by quantitative-PCR (qPCR) using gene-specific primers (Sup. Table S5), as described (Bahar *et al*., 2016). Overall, 17 DEGs from the RNA-seq dataset were tested, using four biological replicates of RNA of OMV- or mock-treated seedlings. Relative expression of the tested genes was compared with ubiquitin expression, using a 7500 Fast real-time PCR machine (Applied Biosystems), as described (Bahar *et al*., 2016).

### Comparing Arabidopsis transcriptional response to MAMPs and to OMVs

Our OMV-induced dataset was compared with available transcriptomes of Arabidopsis challenged with flg22, (Denoux *et al*., 2008) elf26 (Zipfel *et al*., 2006), PGN (Willmann *et al*., 2011), OG (Davidsson *et al*., 2017) and LPS (Livaja *et al*., 2008). For data comparison, GO terms enrichment was performed and the induced GOs were visualized by Venn diagrams as described above.

### Arabidopsis priming experiments

To test the priming effect of *Xcc* OMVs on Arabidopsis infected with *Pseudomonas syringae* pv. *tomato* DC3000 (*Pst*), we followed the procedure described by Zipfel *et al*. (2004). In brief, leaves of 4-5 weeks-old Arabidopsis plants, grown as described above, were infiltrated with 50-100 µL of purified *Xcc* OMVs (30 µg/mL), 1 µM flg22, or water using a needle-less syringe. For each treatment, 5 leaves/plant and three plant replicates were used. *Pst* inoculum was prepared by culturing the bacterium on King’s B medium plates (20 g/L peptone, 1.5 g/L MgSO_4_ x 7 H_2_O, 10 mL/L glycerol and 15 g/L agar) at 28°C for 2-3 days, and then resuspending colonies with water and adjusting the inoculum concentration to 10^5^ CFU/mL. *Pst* inoculum was infiltrated to primed leaves 24 h following priming, using a needle-less syringe (approximately 100 µL were infiltrated to each leaf). Bacterial growth was determined at 0 (1 h post inoculation) and 2 days post inoculation (dpi) by collecting and weighing the inoculated leaves, macerating them in 1 mL of 10 mM MgCl2 and plating 10-fold serial dilutions on King’s B agar plates. The number of CFU on each plate was determined 2 days later and calculate per g leaf.

### *In vitro* bacterial growth assays

To assess the effect of purified *Xcc* OMVs on *Pst* growth *in vitro, Pst* starters were grown in King’s B liquid medium for 24 h at 28°C and then used to inoculate three different cultures containing 12 mL of King’s B medium each, in 50-mL Falcon tubes at a ratio of 1:100. Bacterial cultures were amended with 30 µg/mL OMVs (1:50 or 1:100), or PBS as control, and incubated for 20 h at 28°C. Bacterial growth was measured using a spectrophotometer (Amersham Biosciences) at optical density (OD) of 600 nm over 22 h.

## ACKNOWLEDGEMENTS

The authors wish to thank Prof. Saul Burdman for valuable comments throughout the course of this study and for reviewing drafts of this manuscript.

## FUNGING

This research was supported by the Israel Science Foundation, grant no. 2025/16 to O.B. The funding agency had no role in the design of this study, data collection and analysis, or preparation of the manuscript.

## SUPPLEMENTARY MATERIAL

**Sup. Fig. S1.**
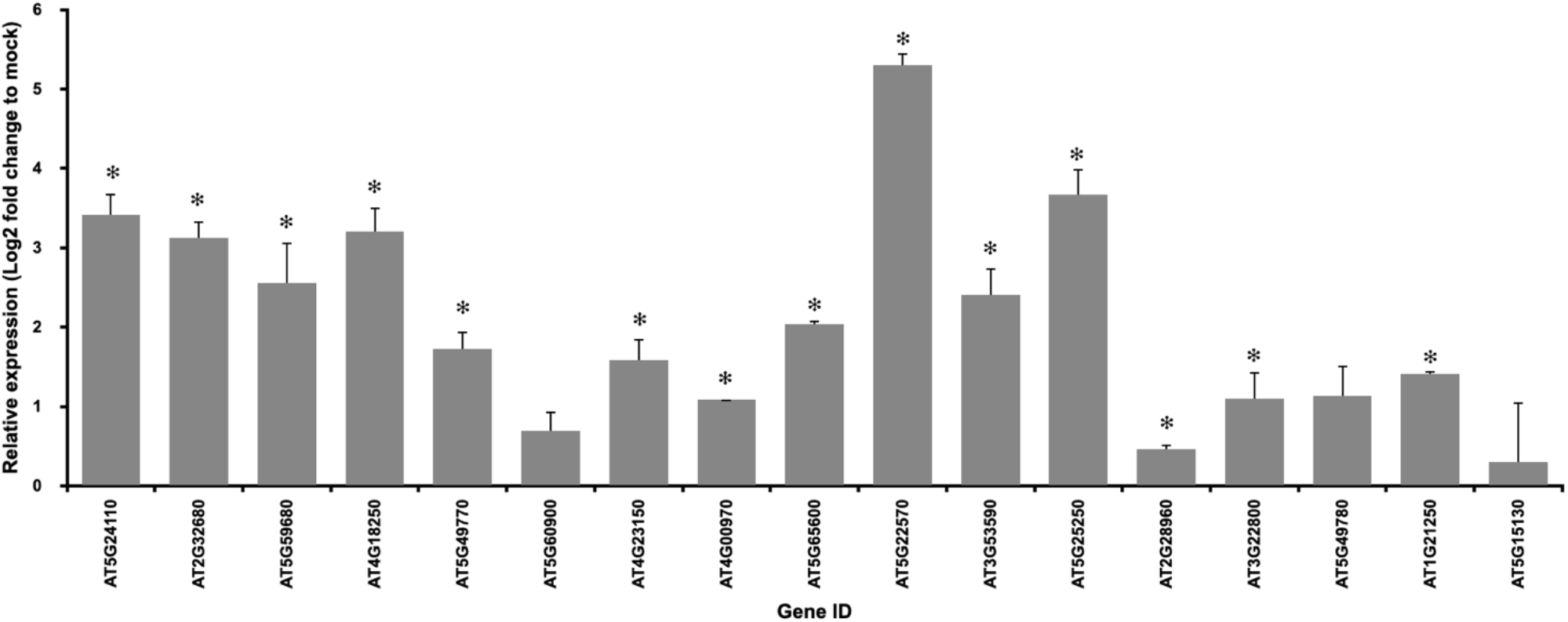
RNA-seq results validation using quantitative-PCR (qPCR). QPCR was performed with 17 differentially expressed genes based on RNA-seq analysis. Specific primers were designed, and the expression of the selected genes was evaluated as described in Material and Methods section. Asterisks indicated statistical significance according to Student t-test (*p*<0.05) compared with mock plants.

**Sup. Fig. S2:**
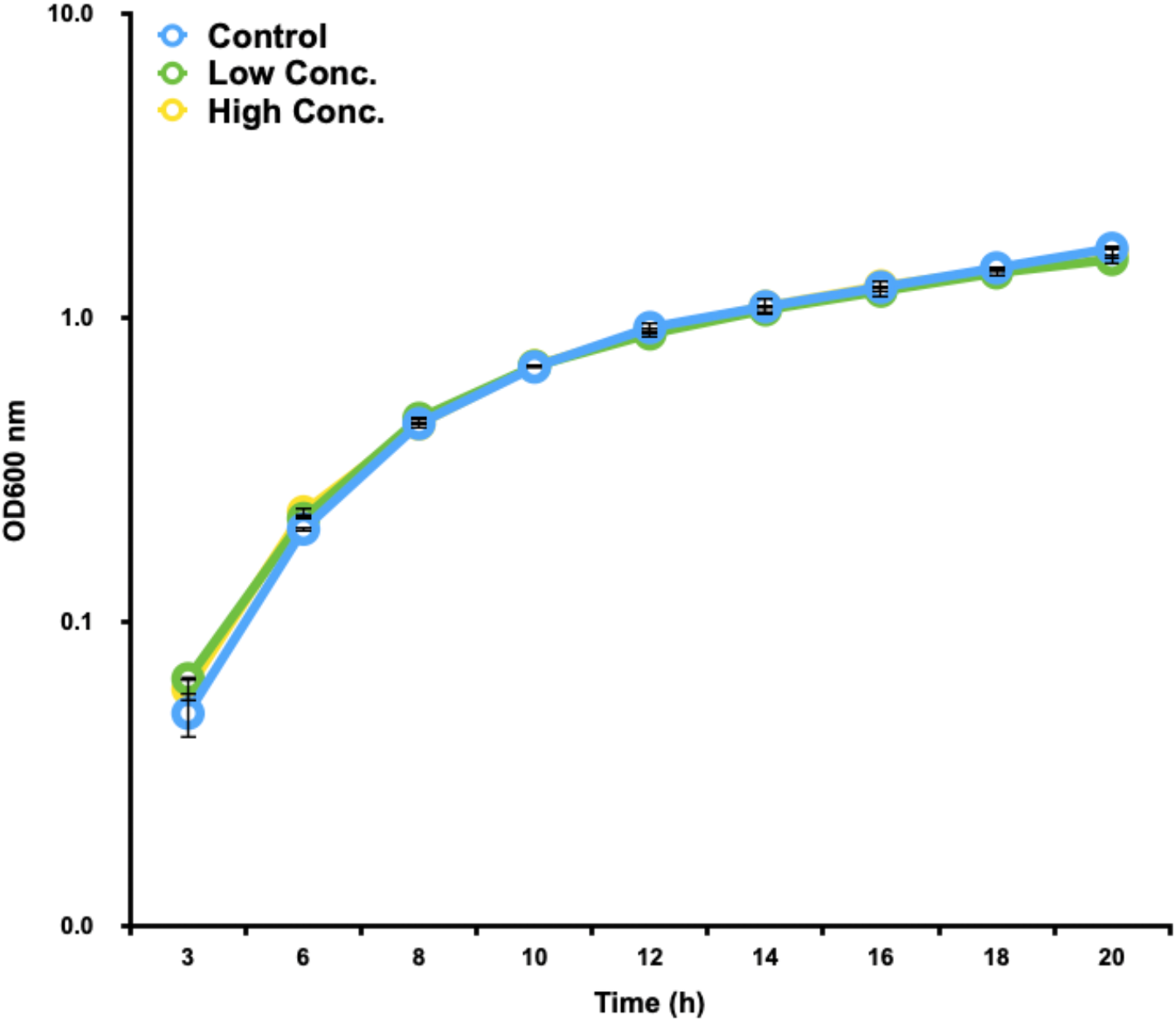
*Pseudomonas syringae* pv. *tomato* (*Pst*) DC3000 growth *in vitro* is not affected by OMVs. *Pst* overnight starters were used to inoculate 50-mL Falcon tubes containing 12 mL of Kings B medium at 1:100 (v:v). *Xcc* OMVs were added to the tubes at two ratios: low (1:100, 0.3 µg/mL), or high (1:50, 0.6 µg/mL). Control tubes contained King’s B medium alone. Cultures were incubated with shaking for 22 h and the OD_600_ of the cultures was monitored every 2 to 3 h. N=3, bars represent standard error of the mean. No statistical differences (ANOVA *p*=0.05) were found at any of the time points tested among the three cultures.

**Sup. Fig. S3:**
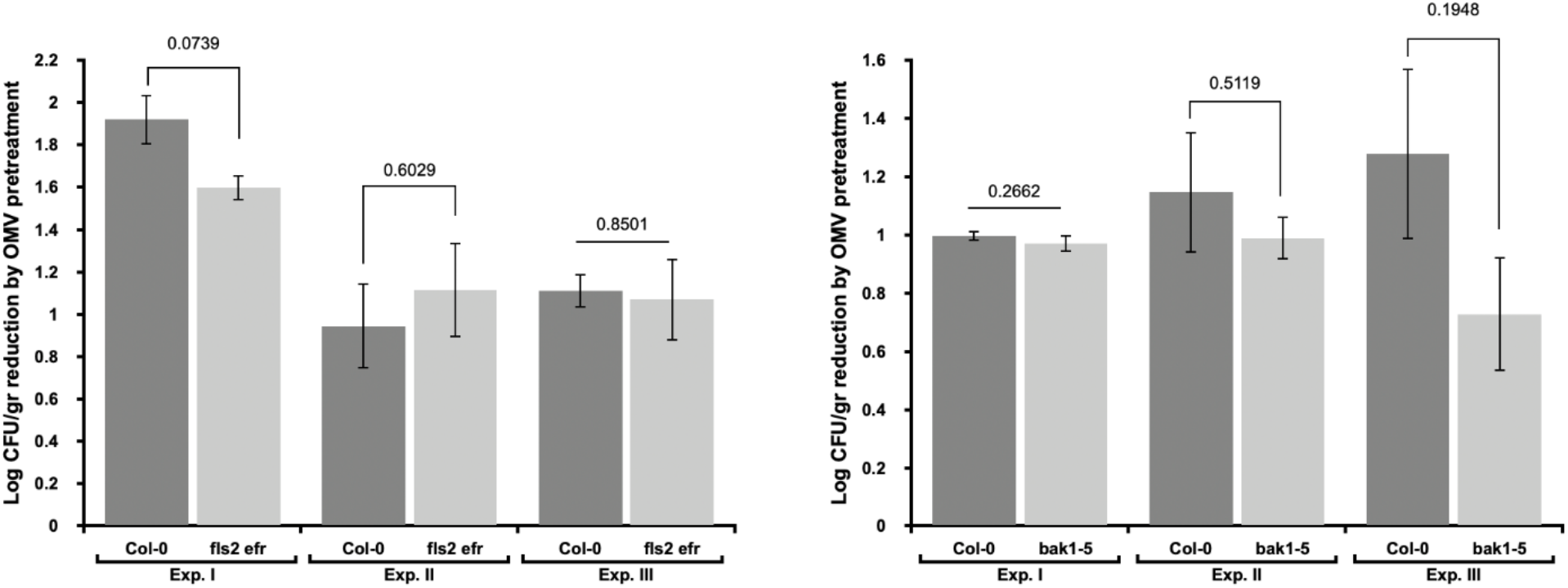
Involvement of EFR, FLS2 and BAK1 receptors in plant priming by OMVs. Arabidopsis Col-0 wild type and *fls2 efr* **(A)** or *bak1-5* **(B)** mutant lines were pretreated with OMVs or water, and then inoculated by *Pst* DC3000, as described in Fig. 5. *Pst* DC3000 cell titer was determined by serial dilution planting and CFU counts 48 h after inoculation (three independent experiments are shown: Exp. I, Exp. II, Exp. III). Bars indicate average *Pst* DC3000 Log fold reduction in Col-0, *fls2 efr* and *bak1-5* lines following OMV pretreatment, compared with untreated plants. No statistical differences were found among both mutant lines and Col-0 in all experiments (*p* values according to Student t-test are shown above graph bars).

## REFERENCES

Ambawat S, Sharma P, Yadav NR, Yadav RC. 2013. MYB transcription factor genes as regulators for plant responses: An overview. Physiology and Molecular Biology of Plants 19: 307–321.

Anders S, Huber W. 2010. Differential expression analysis for sequence count data. Genome Biology 11.

Asai T, Tena G, Plotnikova J, Willmann MR, Chiu WL, Gomez-Gomez L, Boller T, Ausubel FM, Sheen J. 2002. Map kinase signalling cascade in Arabidopsis innate immunity. Nature 415: 977– 983.

Bahar O. 2020. Membrane vesicles from plant pathogenic bacteria and their roles during plantpathogen interactions. In: Kaparakis-Liaskos M, Kufer TA, eds. Bacterial Membrane Vesicles - Biogenesis, Functions and Applications. Springer International Publishing, 119–129.

Bahar O, Mordukhovich G, Luu DD, Schwessinger B, Daudi A, Jehle AK, Felix G, Ronald PC. 2016. Bacterial outer membrane vesicles induce plant immune responses. Molecular Plant-Microbe Interactions 29: 374–384.

Birkenbihl RP, Kracher B, Ross A, Kramer K, Finkemeier I, Somssich IE. 2018. Principles and characteristics of the Arabidopsis WRKY regulatory network during early MAMP-triggered immunity. Plant Journal 96: 487–502.

Bjornson M, Pimprikar P, Nürnberger T, Zipfel C. 2021. The transcriptional landscape of Arabidopsis thaliana pattern-triggered immunity. Nature Plants 7: 579–586.

Bolger AM, Lohse M, Usadel B. 2014. Trimmomatic: a flexible trimmer for Illumina sequence data. Bioinformatics 30: 2114–2120.

Boutrot F, Zipfel C. 2017. Function, discovery, and exploitation of plant pattern recognition receptors for broad-spectrum disease resistance. Annual Review of Phytopathology 55: 257–286.

Chinchilla D, Zipfel C, Robatzek S, Kemmerling B, Nürnberger T, Jones JDG, Felix G, Boller T. 2007. A flagellin-induced complex of the receptor FLS2 and BAK1 initiates plant defence. Nature 448: 497–500.

Cook DE, Mesarich CH, Thomma BPHJ. 2015. Understanding plant immunity as a surveillance system to detect invasion. Annual Review of Phytopathology 53: 541–563.

Couto D, Zipfel C. 2016. Regulation of pattern recognition receptor signalling in plants. Nature Reviews Immunology 16.

Davidsson P, Broberg M, Kariola T, Sipari N, Pirhonen M, Palva ET. 2017. Short oligogalacturonides induce pathogen resistance-associated gene expression in Arabidopsis thaliana. BMC Plant Biology 17: 1–17.

Deatheragea BL, Cooksona BT. 2012. Membrane vesicle release in bacteria, eukaryotes, and archaea: A conserved yet underappreciated aspect of microbial life. Infection and Immunity 80: 1948–1957.

Denoux C, Galletti R, Mammarella N, Gopalan S, Werck D, De Lorenzo G, Ferrari S, Ausubel FM, Dewdney J. 2008. Activation of defense response pathways by OGs and Flg22 elicitors in Arabidopsis seedlings. Molecular Plant 1: 423–445.

Dow M, Newman M-A, von Roepenack E. 2000. The induction and modulation of plant defense responses by bacterial lipopolysaccharides. Annual Reviews of Phytopathology 38: 241–261.

Du Z, Zhou X, Ling Y, Zhang Z, Su Z. 2010. agriGO: A GO analysis toolkit for the agricultural community. Nucleic Acids Research 38: W64–W70.

Ellis TN, Kuehn MJ. 2010. Virulence and immunomodulatory roles of bacterial outer membrane vesicles. Microbiology and Molecular Biology Reviews 74: 81–94.

Ellis TN, Leiman SA, Kuehn MJ. 2010. Naturally produced outer membrane vesicles from Pseudomonas aeruginosa elicit a potent innate immune response via combined sensing of both lipopolysaccharide and protein components. Infection and Immunity 78: 3822–3831.

Erbs G, Silipo A, Aslam S, de Castro C, Liparoti V, Flagiello A, Pucci P, Lanzetta R, Parrilli M, Molinaro A, et al. 2008. Peptidoglycan and muropeptides from pathogens Agrobacterium and Xanthomonas elicit plant innate immunity: structure and activity. Chemistry and Biology 15: 438– 448.

Felix G, Duran JD, Volko S, Boller T. 1999. Plants have a sensitive perception system for the most conserved domain of bacterial flagellin. Plant Journal 18: 265–276.

Fesel PH, Zuccaro A. 2016. β-glucan: Crucial component of the fungal cell wall and elusive MAMP in plants. Fungal Genetics and Biology 90: 53–60.

Fulsundar S, Harms K, Flaten GE, Johnsen PJ, Chopade BA, Nielsen KM. 2014. Gene transfer potential of outer membrane vesicles of Acinetobacter baylyi and effects of stress on vesiculation. Applied and environmental microbiology 80: 3469–83.

Gómez-Gómez L, Boller T. 2000. FLS2: An LRR receptor-like kinase involved in the perception of the bacterial elicitor flagellin in Arabidopsis. Molecular Cell 5: 1003–1011.

Gurung M, Moon DC, Choi CW, Lee JH, Bae YC, Kim J, Lee YC, Seol SY, Cho DT, Kim SI, Lee JC. 2011. Staphylococcus aureus produces membrane-derived vesicles that induce host cell death. PLoS ONE, 6(11).

Gust AA, Biswas R, Lenz HD, Rauhut T, Ranf S, Kemmerling B, Gotz F, Glawischnig E, Lee J, Felix G, et al. 2007. Bacteria-derived peptidoglycans constitute pathogen-associated molecular patterns triggering innate immunity in Arabidopsis. Journal of Biological Chemistry 282: 32338– 32348.

Ionescu M, Zaini PA, Baccari C, Tran S, da Silva AM, Lindow SE. 2014. Xylella fastidiosa outer membrane vesicles modulate plant colonization by blocking attachment to surfaces. Proceedings of the National Academy of Sciences of the United States of America 111: E3910–E3918.

Janda M, Ludwig C, Rybak K, Meng C, Stigliano E, Botzenhardt L, Szulc B, Sklenar J, Menke FLH, Malone JG, et al. 2021. Biophysical and proteomic analyses suggest functions of Pseudomonas syringae pv tomato DC3000 extracellular vesicles in bacterial growth during plant infection. bioRxiv: 2021.02.08.430144.

Jin JS, Kwon SO, Moon DC, Gurung M, Lee JH, Kim S, Lee JC. 2011. Acinetobacter baumannii secretes cytotoxic outer membrane protein a via outer membrane vesicles. PLoS ONE, 6(2).

Jung HW, Tschaplinski TJ, Wang L, Glazebrook J, Greenberg JT. 2009. Priming in systemic plant immunity. Science 324: 89–91.

Kadurugamuwa JL, Beveridge TJ. 1996. Bacteriolytic effect of membrane vesicles from Pseudomonas aeruginosa on other bacteria including pathogens: conceptually new antibiotics. Joutnal of Bacteriology 178: 2767–2774.

Katsir L, Bahar O. 2017. Bacterial outer membrane vesicles at the plant–pathogen interface. PLoS Pathogens 13: 1–6.

Kemmerling B, Halter T, Mazzotta S, Mosher S, Nürnberger T. 2011. A genome-wide survey for Arabidopsis leucine-rich repeat receptor kinases implicated in plant immunity. Frontiers in Plant Science 2: 88.

Kuehn MJ, Kesty NC. 2005. Bacterial outer membrane vesicles and the host-pathogen interaction. Genes and Development 19: 2645–2655.

Kunsmann L, Rüter C, Bauwens A, Greune L, Glüder M, Kemper B, Fruth A, Wai SN, He X, Lloubes R, et al. 2015. Virulence from vesicles: Novel mechanisms of host cell injury by Escherichia coli O104:H4 outbreak strain. Scientific Reports 5: 13252.

Lamesch P, Berardini TZ, Li D, Swarbreck D, Wilks C, Sasidharan R, … Huala E. 2012. The Arabidopsis Information Resource (TAIR): Improved gene annotation and new tools. Nucleic Acids Research, 40: 1202–1210.

Langmead, B, Salzberg SL. 2012. Fast gapped-read alignment with Bowtie 2. Nature Methods: 9:357–359.

Li B, Dewey CN. 2011. RSEM: accurate transcript quantification from RNA-Seq data with or without a reference genome. BMC Bioinformatics 12: 323.

Livaja M, Zeidler D, von Rad U, Durner J. 2008. Transcriptional responses of Arabidopsis thaliana to the bacteria-derived PAMPs harpin and lipopolysaccharide. Immunobiology 213: 161–171.

Love MI, Huber W, Anders S. 2014. Moderated estimation of fold change and dispersion for RNA-seq data with DESeq2. Genome Biology 15: 1–21.

MacDonald IA, Kuehn MJ. 2013. Stress-induced outer membrane vesicle production by Pseudomonas aeruginosa. Journal of bacteriology 195: 2971–2981.

Manning AJ, Kuehn MJ. 2011. Contribution of bacterial outer membrane vesicles to innate bacterial defense. BMC Microbiology 11.

Mashburn LM, Whiteley M. 2005. Membrane vesicles traffic signals and facilitate group activities in a prokaryote. Nature 437: 422–425.

McMillan HM, Zebell SG, Ristaino JB, Dong X, Kuehn MJ. 2021. Protective plant immune responses are elicited by bacterial outer membrane vesicles. Cell Reports 34: 108645.

McMillan HM, Kuehn MJ. 2021. The extracellular vesicle generation paradox: a bacterial point of view. The EMBO Journal, 1–23.

Mordukhovich G, Bahar O. 2017. Isolation of outer membrane vesicles from phytopathogenic Xanthomonas campestris pv. campestris. Bio-Protocol 7: 1–13.

Mott GA, Thakur S, Smakowska E, Wang PW, Belkhadir Y, Desveaux D, Guttman DS. 2016. Genomic screens identify a new phytobacterial microbe-associated molecular pattern and the cognate Arabidopsis receptor-like kinase that mediates its immune elicitation. Genome Biology 17: 98.

Nekrasov V, Li J, Batoux M, Roux M, Chu ZH, Lacombe S, Rougon A, Bittel P, Kiss-Papp M, Chinchilla D, et al. 2009. Control of the pattern-recognition receptor EFR by an ER protein complex in plant immunity. EMBO J. 28: 3428–3438.

Porter K, Shimono M, Tian M, Day B. 2012. Arabidopsis actin-depolymerizing factor-4 links pathogen perception, defense activation and transcription to cytoskeletal dynamics. PLoS Pathogens 8.

Ranf S, Scheel D, Lee J. 2016. Challenges in the identification of microbe-associated molecular patterns in plant and animal innate immunity: a case study with bacterial lipopolysaccharide. Molecular plant pathology 17: 1165–1169.

Raposo G, Stahl PD. 2019. Extracellular vesicles: a new communication paradigm? Nature Reviews Molecular Cell Biology 20: 509–510.

Rushton PJ, Somssich I. E, Ringler P, Shen Q. 2010. WRKY transcription factors. Trends in Plant Science 15: 247–258.

Schooling SR, Beveridge TJ. 2006. Membrane vesicles: An overlooked component of the matrices of biofilms. Journal of Bacteriology 188: 5945–5957.

Schwechheimer C, Kuehn MJ. 2015. Outer-membrane vesicles from Gram-negative bacteria: biogenesis and functions. Nature Reviews Microbiology 13: 605–619.

Schwessinger B, Roux M, Kadota Y, Ntoukakis V, Sklenar J, Jones A, Zipfel C. 2011. Phosphorylation-dependent differential regulation of plant growth, cell death, and innate immunity by the regulatory receptor-like kinase BAK1. PLoS genetics 7: e1002046.

Silipo A, Molinaro A, Sturiale L, Dow JM, Erbs G, Lanzetta R, Newman MA, Parrilli M. 2005. The elicitation of plant innate immunity by lipooligosaccharide of Xanthomonas campestris. J Biol Chem 280: 33660–33668.

Silipo A, Sturiale L, Garozzo D, Erbs G, Jensen TT, Lanzetta R, Dow JM, Parrilli M, Newman MA, Molinaro A. 2008. The acylation and phosphorylation pattern of lipid A from Xanthomonas campestris strongly influence its ability to trigger the innate immune response in Arabidopsis. ChemBioChem 9: 896–904.

Tian T, Liu Y, Yan H, You Q, Yi X, Du Z, Xu W, Su Z. 2017. AgriGO v2.0: A GO analysis toolkit for the agricultural community, 2017 update. Nucleic Acids Research 45: W122–W129.

Tran TM, Chng C, Pu X, Ma Z. 2021. Potentiation of plant defense by bacterial outer membrane vesicles is mediated by membrane nanodomains. The Plant Cell, 1–24.

Tsuda K, Somssich IE. 2015. Transcriptional networks in plant immunity. New Phytologist 206: 932–947.

Velimirov B, Ranftler C. 2018. Unexpected aspects in the dynamics of horizontal gene transfer of prokaryotes: the impact of outer membrane vesicles. Wiener Medizinische Wochenschrift 168: 307– 313.

Wang G, Ellendorff U, Kemp B, Mansfield JW, Forsyth A, Mitchell K, Bastas K, Liu C-M, Woods-Tor A, Zipfel C, et al. 2008. A genome-wide functional investigation into the roles of receptor-like proteins in Arabidopsis. Plant Physiology 147: 503–517.

Willmann R, Lajunen HM, Erbs G, Newman MA, Kolb D, Tsuda K, Katagiri F, Fliegmann J, Bono JJ, Cullimore J V, et al. 2011. Arabidopsis lysin-motif proteins LYM1 LYM3 CERK1 mediate bacterial peptidoglycan sensing and immunity to bacterial infection. Proceedings of the National Academy of Sciences of the United States of America 108: 19824–19829.

Zipfel C, Kunze G, Chinchilla D, Caniard A, Jones JD, Boller T, Felix G. 2006. Perception of the bacterial PAMP EF-Tu by the receptor EFR restricts Agrobacterium-mediated transformation. Cell 125: 749–760.

Zipfel C, Robatzek S, Navarro L, Oakeley EJ, Jones JDG, Felix G, Boller T. 2004. Bacterial disease resistance in Arabidopsis through flagellin perception. Nature 428: 764–767.

